# Bridging functional annotation gaps in non-model plant genes with AlphaFold, DeepFRI and small molecule docking

**DOI:** 10.1101/2021.12.22.473925

**Authors:** Georgie Stephan, Benjamin Dugdale, Pradeep Deo, Rob Harding, James Dale, Paul Visendi

## Abstract

**Background:** Functional annotation assigns descriptive biological meaning to genetic sequences. Limited availability of manually curated or experimentally validated plant genes from a diverse range of taxa poses a significant challenge for functional annotation in non-model organisms. Accurate computational approaches are required. We argue that recent breakthroughs in deep learning have the potential to not only narrow the functional annotation gap between non-model and model plant organisms, but also annotate and reveal novel functions even for genes with no homologs in public databases.

**Results:** Deep learning models were applied to functionally annotate a set of previously published differentially expressed genes. Predicted protein structures and functional annotations were generated using the AlphaFold protein structure and DeepFRI protein language inference models respectively. The resulting structures and functional annotations were validated using small molecule docking experiments. DeepFRI and AlphaFold models not only correctly annotated differentially expressed genes, but also revealed detailed mechanisms involving protein-protein interactions.

**Conclusions:** Deep learning models are capable of inferring novel functions and achieving high accuracy in functional annotation. Their increased use in plant research will result in major improvements in annotations for non-model plants that are underrepresented in genome databases. We illustrate how integrating protein structure prediction, functional residue prediction, and small molecule docking can infer plausible protein-protein interactions and yield additional mechanistic insights. This approach will aid in the selection of candidate genes for further study from differential expression studies that generate large gene lists.

## Background

Current methods for genome functional annotation at genome scale in genomics rely on homology transfer and inference from experimentally or manually curated and functionally annotated genes in public databases such as the Protein Data Bank (PDB) and UniProtKB/Swiss-Prot. Only 0.5% of the roughly 100 million sequences deposited in UniProtKB are manually curated ^1^. Similarly, just 624 (3.9%) of the 15,897 experimentally determined structures in the PDB (as of July 2021) are from plants. Most functional annotations for plants are based on *Arabidopsis* as the model organism. This introduces a significant annotation bias, given the fact that there are about 400,000 plant species ^2^, the vast majority of which have not been sequenced.

Although the UniProtKB/Swiss-Prot manually curated protein databases contain a higher proportion of plant sequences (21%) (UniProtKB/Swiss-Prot release 2021 03 on June 2021), these are based on automated homology-based assignments with additional manual curation from studies involving mostly model organisms. Manual curation entails the assessment and integration of functional evidence from scientific literature by a human expert. Deep learning approaches for protein structure and function prediction have been shown to generate more accurate functional annotations, even for novel sequences with no homologs in the PDB or UniProtKB/Swiss-Prot databases ^3^. This is important considering the lower representation of plant-specific sequences in the PDB and UniProtKB/Swiss-Prot.

The biological activity of a gene is determined by the unique three-dimensional structure of the protein it encodes and the availability or identification and functional annotation of protein-coding genes in an organism serves as the foundation for any genomic analysis. The rapid accumulation of new genomic datasets generated by large-scale sequencing projects necessitates application of automated functional annotation approaches that can both annotate large datasets of genomic sequences efficiently but are also capable of annotating novel sequences.

Experimental determination of protein structures or functionally validating all genes in a newly sequenced non-model organism is difficult and generally intractable. Decades of theoretical work on predicting protein structures and functions directly from amino acid sequences using deep learning have finally come to fruition thanks to current state-of-the-art deep learning technologies. Deep learning algorithms can produce results comparable to those of empirical laboratory experiments and provide a strong rationale for adopting these methods more widely to address the functional annotation gaps inherent in non-model plant sequences.

Although experimental protein structure determination methods have improved in accuracy in recent years, they remain time-consuming and expensive ^4^. Until now, predicting three-dimensional protein structures based solely on amino acid sequences has been an ongoing bioinformatics challenge. In 2020-2021, cutting-edge deep learning algorithms such as DeepMind’s AlphaFold ^5,6^ and the Baker Lab’s RoseTTAFold ^7^ demonstrated that predicted protein structures are accurate and can be indistinguishable or better than experimentally determined structures when compared to gold standard methods such as X-ray crystallography and cryogenic electron microscopy (cryo-EM) ^8^. AlphaFold is based on a convolutional neural network trained on experimentally determined protein structures and their amino acid sequences from the PDB. AlphaFold accurately models distances between amino acid residues in three-dimensional space based on their spatial orientation in protein structures. The models when applied allow accurate prediction of a protein’s three-dimensional structure directly from its amino acid sequence. AlphaFold was able to predict previously unknown folds of proteins without sequence or structural homologs in the PDB at high atomic accuracy. Because protein structures exhibit three to ten times more evolutionary conservation in topology or folds than the underlying linear amino acid sequences ^9^, deep learning models can learn evolutionary conserved topologies and accurately predict protein structures even of previously uncharacterised proteins.

Based on a similar architecture to AlphaFold, RoseTTAFold ^7^ also predicts three-dimensional structures directly from amino acid sequences with accuracies approaching those of AlphaFold. Unlike AlphaFold, RoseTTAFold uses a three-network track model where linear amino acid sequences (1D) on one track are coupled with their respective distances based on a 2D map on a second track and finally atomic coordinates are refined in 3D on the third track. RoseTTAFold results further confirmed that accurate models for proteins with no or partial homologs in the PDB can be predicted. The bacterial surface layer protein (SLP) and the Lrbp protein from the fungus *Phanerochaete chrysosporium* are such examples. RoseTTAFold was able to generate accurate structures of *G* protein coupled receptors in both active and inactive states. This suggests that deep learning models can capture multiple protein conformations and thus can provide new insights into a protein’s function. Most significant is that both RoseTTAFold and AlphaFold can generate structures of protein-protein complexes directly from sequence information without the need for protein-protein docking ^7,10^. In addition to functional annotation of novel genes, these findings will advance protein-protein interaction studies.

The structural bioinformatics community runs an event, Critical Assessment of Structure Prediction (CASP), every two years that benchmarks the accuracy of protein structure prediction methods against a common set of targets. The event is open to any research group involved in protein structure prediction. Independent researchers contribute newly determined crystal structures to CASP prior to public release on the PDB. The protein sequences for these structures are subsequently made available to CASP contenders, who without knowledge of the crystal structures, compete to accurately predict the protein structures. Submitted predictions are compared to crystal structures and ranked using a global distance test (GDT) score ranging from 0 to 100. A score of 100 indicates that the predicted structure is identical to the crystal structure. For an unbiased assessment, the protein targets are grouped into template-based modelling categories for proteins with considerable sequence homologs in the PDB and the free modelling categories for proteins with no structural homologs in the PDB.

In the 2020 CASP challenge, AlphaFold not only achieved accuracies with median GDT scores of 92.4 but was able to solve structures for proteins with no homologs in the PDB with median GDT scores of 87 ^4^. GDT scores above 90 are considered accurate enough to infer biological mechanisms. This was a significant finding that demonstrated accurate structures of novel proteins can now be predicted and provided a major step towards bridging the annotation gaps for less characterized plants and non-model species. AlphaFold has been further demonstrated to accurately predict protein-protein complexes ^11‒13^ and has already been used to generate protein models of more than 20 species accessible at the AlphaFold Protein Structure Database (https://alphafold.ebi.ac.uk).

Protein language models now enable residue level, site-specific functional annotations, allowing unprecedented resolution and insight into protein function. Blind prediction benchmarks set by the Critical Assessment of Functional Annotation (CAFA) challenge shows that application of deep learning methods to predict site-specific functions continues to outperform homology-based methods such as BLAST ^14^. Most protein function prediction methods that use protein sequences as inputs use convolutional neural networks (CNNs). CNNs search for spatial patterns in a database of protein sequences such as the PDB. Extracted patterns are then used as feature inputs into the CNN ^15^.

Graph convolutional networks (GCNs), a variation of CNNs, have increasingly been applied for building protein function prediction models ^16^. Deep Functional Residue Identification (DeepFRI), a recent model based on GCNs ^1^ showed predictions of functional residue gene ontology (GO) terms are robust to errors in protein structure models. To predict residue functions based on the position of a residue in an amino acid sequence, DeepFRI was trained on a dataset of 10 million sequences from the protein families (Pfam) database ^17^.

For accurate functional annotation transfer using homology-based approaches, an appropriate sequence alignment must be generated. Depending on the parameter settings applied during homology searches, sequence alignments for the same dataset can yield differing results. For example, sequence match lengths, sequence identity thresholds, and the complexity of the target to which sequences are aligned are all factors to consider. When combined with protein structures, DeepFRI identifies and annotates functionally important residues that may be distal in a linear amino acid sequence but proximal in the three-dimensional structure, outperforming homology-based methods ^1^. This approach negates the need to perform computationally intensive homology searches using arbitrarily set homology match parameters.

We show using several genes how molecular and enzymatic functions can be predicted by first predicting protein structures, identifying functional residues in these structures, and validating catalytic sites using small molecule docking experiments. As an example, we use AlphaFold to predict protein structures and DeepFRI to generate functional annotations at the residue level for 19 differentially expressed genes from a published differential expression study in bananas ^18^, a non-model plant. In this study, differentially expressed genes were analysed from leaf tissues of the susceptible Cavendish banana (*Musa acuminata*) cultivar Williams and a resistant cultivar Calcutta 4 (*Musa acuminata* ssp. *burmannicoides*), after inoculation with *Pseudocercospora fijiensis*, the causative agent for black Sigatoka. The lowest free energy conformation AlphaFold structure predictions for each of the 19 genes was taken as input in to DeepFRI. DeepFRI uses a gradient-weighted Class Activation Map (grad-CAM) to identify residues associated with the predicted GO term probabilities ^19^. We then endeavoured to infer protein-protein interactions from the generated functional annotations in the context of the differential expression study. Finally, we predicted structures of protein-protein complexes for several genes and validated these using *in silico* small molecule docking.

## Results

### AlphaFold and RosettaFold comparisons

Using TM-align ^20^, a comparison of AlphaFold and RosettaFold predicted structures for the 19 protein sequences revealed that both methods predicted similar folds (avg TM-score of 0.5). Two protein structures with a TM-Score of 0.2 are considered to have distinct folds, whereas those with a score greater than or equal to 0.5 are considered to have the same fold.

Both AlphaFold and RoseTTAFold rank the accuracy of models based on predicted Local Distance Difference Test (pLDDT) scores ^21,22^ with AlphaFold scaling pLDDT values between 0 to 100 and RoseTTAFold between 0 to 1. To get an estimate of the comparative accuracy of AlphaFold and RoseTTAFold models of the 19 protein sequences, both AlphaFold and RoseTTAFold pLDDT values (Table 1) were re-scaled to values between 0 and 1. We used a similar approach to Visendi *et al*., (2016)^23^ to statistically compare the performance of different algorithms on the same dataset. In this case, a Wilcoxon signed-rank test was applied as the pLDDT scores were not normally distributed. AlphaFold scaled pLDDT values were significantly higher (*P* value < 0.008) than RoseTTAFold values (Fig 1a). AlphaFold scaled pLDDT scores were positively correlated with TM-scores (Spearman-r of 0.70 with *P* value < 0.0007) (Fig 1b). RoseTTAFold pLDDT scores were also positively correlated with TM-scores (Spearman-r of 0.77 with *P* value < 0.0001) (Fig 1c).

**Table 1:** TM-Scores, AlphaFold and RosettaFold raw and scaled pLDDT scores.

| Protein | TM-score | protein aa length | AlphaFold pLDDT | AlphaFold | RosettaFold pLDDT | RosettaFold |
| --- | --- | --- | --- | --- | --- | --- |
| ZAT10 | 0.15907 | 269 | 63.32 | 0.291701593 | 0.38 | 0 |
| TIFY38 | 0.26009 | 228 | 60.22 | 0.226739313 | 0.45 | 0.134615385 |
| RD22 | 0.72675 | 351 | 74.44 | 0.524727578 | 0.69 | 0.596153846 |
| PYL8 | 0.80608 | 227 | 83.85 | 0.721919531 | 0.73 | 0.673076923 |
| PR1 | 0.95558 | 160 | 97.12 | 1 | 0.9 | 1 |
| OPR1 | 0.9663 | 365 | 94.54 | 0.945934619 | 0.86 | 0.923076923 |
| MYC2 | 0.31625 | 696 | 66.75 | 0.363579212 | 0.4 | 0.038461538 |
| MYB | 0.39231 | 275 | 64.61 | 0.318734283 | 0.57 | 0.365384615 |
| LOX8 | 0.27711 | 63 | 49.4 | 0 | 0.55 | 0.326923077 |
| IAA30 | 0.43575 | 247 | 62.5 | 0.274518022 | 0.45 | 0.134615385 |
| IAA27 | 0.36269 | 322 | 58.01 | 0.180427494 | 0.39 | 0.019230769 |
| ERF105 | 0.26559 | 236 | 65.04 | 0.32774518 | 0.45 | 0.134615385 |
| ERD15 | 0.23983 | 178 | 63.2 | 0.289186924 | 0.44 | 0.115384615 |
| EIN3 | 0.27432 | 528 | 66.33 | 0.354777871 | 0.45 | 0.134615385 |
| DREB1D | 0.46805 | 225 | 65.73 | 0.342204526 | 0.53 | 0.288461538 |
| COI1 | 0.96392 | 588 | 93.63 | 0.926865046 | 0.83 | 0.865384615 |
| ASR3 | 0.33302 | 340 | 70.63 | 0.44488684 | 0.44 | 0.115384615 |
| ARP1 | 0.30355 | 280 | 59.16 | 0.204526404 | 0.42 | 0.076923077 |
| ARF7 | 0.53287 | 653 | 69.39 | 0.418901928 | 0.56 | 0.346153846 |

**Fig 1:**
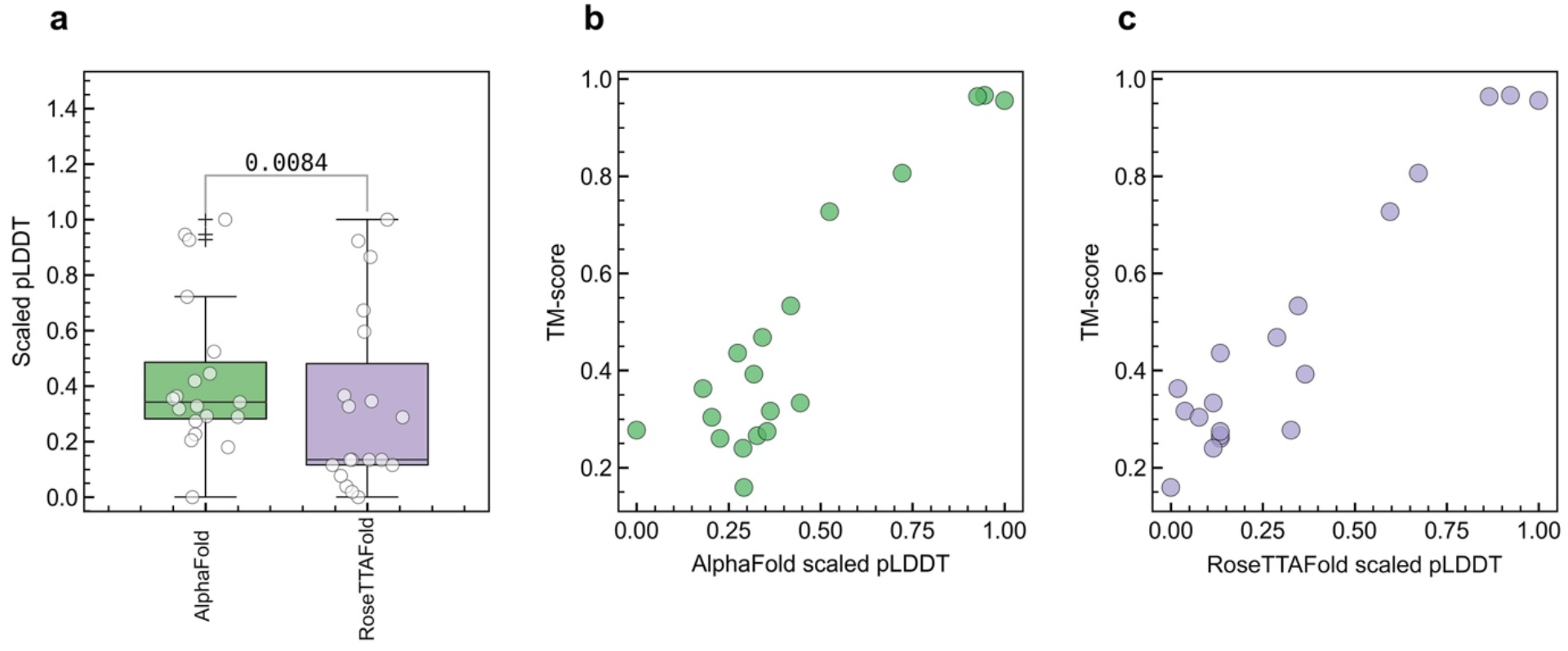
***a)*** AlphaFold vs RosettaFold scalled pLDDT scores AlphaFold has significantly higher pLDDT scores than RosettaFold for the 19 predicted protein structures. ***b, c*)** A high correlation between TM-scores and AlphaFold and RosettaFold pLDDT scores indicates both methods predict similar folds.

Taken together, these results suggest that while AlphaFold and RosettaFold predict similar folds, AlphaFold significantly outperforms RosettaFold in accuracy as has been previously reported. We thus proceeded to generate functional annotations using AlphaFold structure predictions (Fig 2).

**Fig 2:**
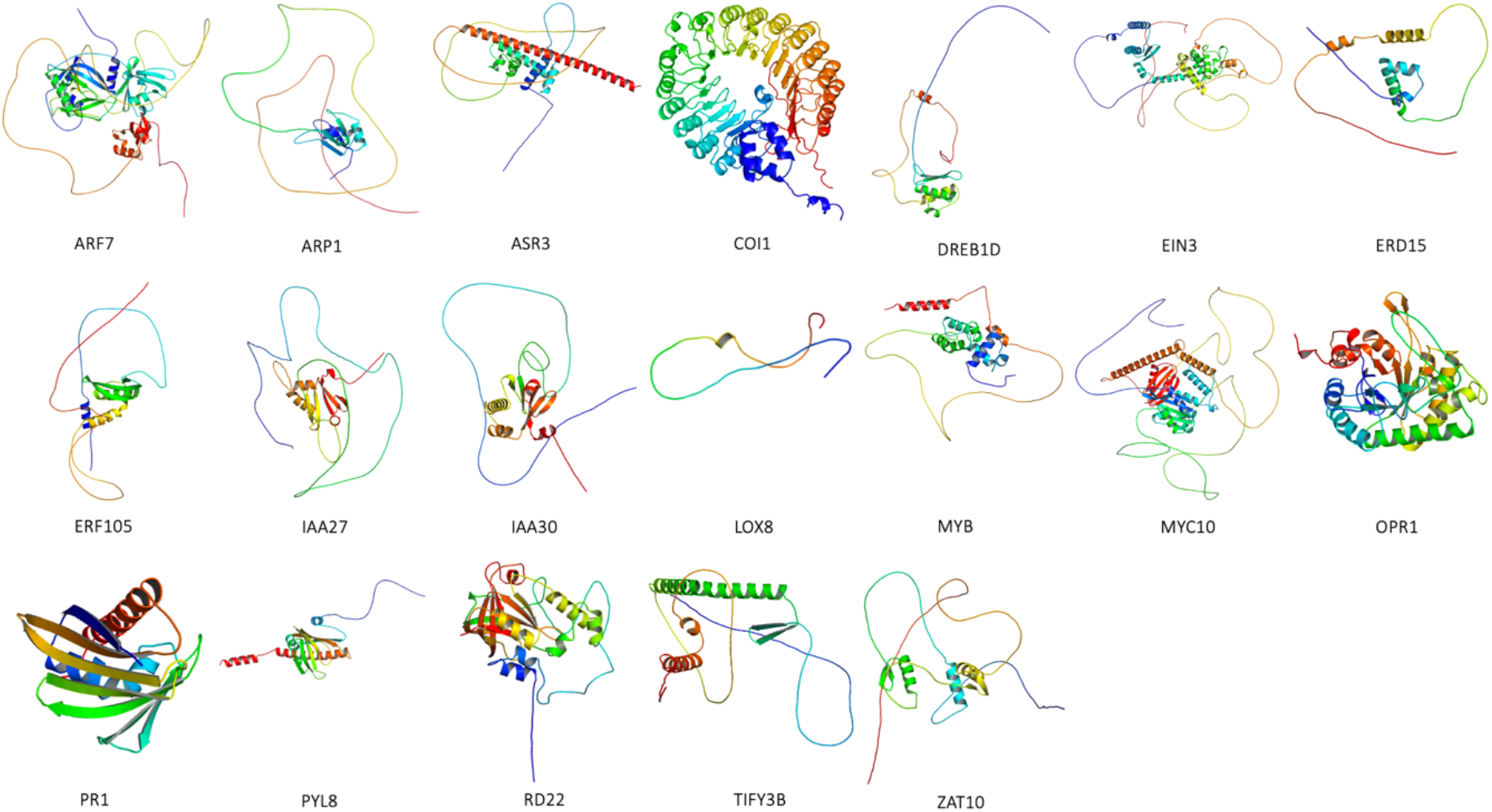
AlphaFold predicted structures of 19 differentially expressed genes reported by Rodriquez *et al*., *2020*. Structures were visualized in Pymol.

### Functional annotations

Using AlphaFold protein structure predictions, DeepFRI models were used to predict molecular function (MF) and enzyme commission (EC) descriptions. DeepFRI calculated probability scores for functional residue catalytic and binding activity ranging from 0.1 to 0.9, with a mean of 0.2 (Table S1). A total of 108 MF GO terms and 68 EC numbers were assigned to the 19 AlphaFold protein models. Each gene had at least one GO term or EC number assigned. Lipoxygenase 9 *LOX8* and *TIFY3B* genes did not have a MF term assigned while dehydration-responsive element-binding 1D (*DREB1D)*, early responsive to dehydration 15 protein (*ERD15)*, auxin response protein 27 (*IAA27)*, auxin response protein 30 (*IAA30)* and dehydration response protein RD22 (*RD22)* had no EC numbers predicted (Table S1).

GO terms and EC numbers are organised hierarchically, with parent GO terms having broad descriptive ontologies and deeper child terms having more specific terms. DeepFRI reports lower-level GO terms. In cases where both parent and child terms were reported for a gene, resulting in annotation redundancy, the deepest MF or EC child term was used for further analysis.

GO terms and EC numbers are organised hierarchically, with parent GO terms having broad descriptive ontologies and deeper child terms having more specific terms. DeepFRI reports lower-level GO terms. In cases where both parent and child terms were reported for a gene, resulting in annotation redundancy, the deepest MF or EC child term was used for further analysis. Using these criteria, we observed that the transcription factor MYC2 (*MYC2)* was the only gene that shared EC assignments with three other genes coronatine-insensitive protein (*COI1)*, MYB family transcription factor *(MYB) and* ET-insensitive 3 (*EIN3*) (Fig 3, Table S1). These were *MYC2* and *COI1* annotated as both encoding *RNA helicase* and *DNA helicase* activity (EC: 3.6.4.13 and EC: 3.6.4.12). *MYC2* and *EIN3* were annotated as sharing *ubiquitinyl hydrolase 1* activity (EC: 3.4.19.12) while *MYC2* and *MYB* shared *histone acetyltransferase* activity (EC: 2.3.1.48). Plots of class activation map probabilities per residue on *MYC2* indicated that unique residue groupings (peaks on Fig 3 a, b, c, d) are predicted to encode *RNA helicase, DNA helicase, ubiquitinyl hydrolase 1* and *histone acetyltransferase* activity on *MYC2*, respectively. This suggests that *MYC2* has multiple functions, may physically interact, or is involved in regulation of or is regulated by multiple gene products. This also suggested that *MYC2* may function as part of a complex with other gene products. These observations are consistent with published findings that *MYC2* is a transcription factor that regulates multiple defence response pathways including the jasmonic acid (JA), salicylic acid (SA), gibberellic acid (GA), abscisic acid (ABA), and indole-3-acetic acid (IAA) pathways ^24,25^. In addition, *MYC2* has been shown to recruit nucleosome remodelling proteins, specifically *histone acetyltransferases* for promoter binding ^26^ further supporting predictions of shared *histone acetyltransferase* activity between *MYC2* and *MYB*.

**Fig 3:**
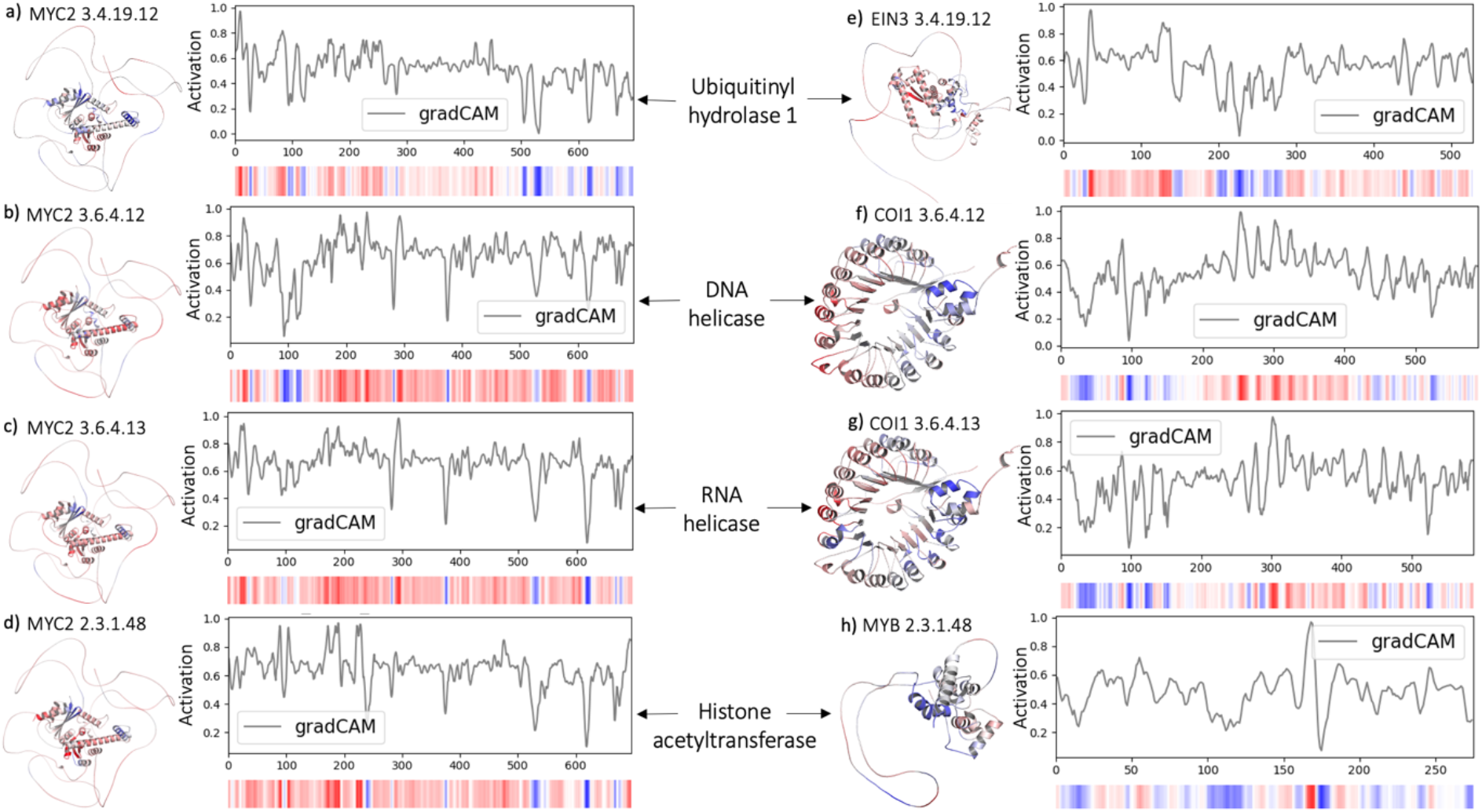
Predicted functional annotations of *MYC2* (***a, b, c, d*)**, *EIN3* (*e*), *COI1* (***f, g***), and *MYB* (***h***) structures, highlighting active residues and the matching gradient-weighted class activation map (grad-CAM) profiles per residue. The left panel represents four predicted MYC2 functions, whereas the right panel represents the corresponding shared functions predicted for EIN3, COI1, and MYB. Residues on the structures and the grad-CAM profile heatmaps are coloured using a divergent red, white, blue gradient. The more active residues are coloured red, whereas the less active residues are highlighted in blue.

While the regulatory role of *MYC2* is well established, the proteins responsible for its phosphorylation remain unclear; see Kazan and Manners, (2013)^24^ for a comprehensive review. We explored whether *EIN3, COI1*, and *MYB* EC assignments shared with *MYC2* could be relied on as an indicator of the likely phosphate donor. *COI1* (EC: 3.6.4.13/12) is predicted as an *anhydride hydrolase* that acts on adenosine triphosphate (ATP) with release of phosphates and adenosine diphosphate (ADP). *EIN3* is predicted to be a peptidase, specifically a *ubiquitinyl hydrolase* (EC: 3.4.19.12). Ubiquitination is key in the regulation of phosphate acquisition and utilization by plants ^27^. *MYB* was predicted to be an acetyltransferase (EC: 2.3.1.48, *histone acetyltransferase*). Based on these functional annotations, it likely that *COI1* is involved in the phosphorylation and *MYB* acetylation. Based on the shared functional residue annotations, we investigated two post-translational protein modifications processes involving *MYC2*, phosphorylation and acetylation, using predicted structures of protein complexes of *MYC2-COI1* (pLDDT 69.15, FigS1) and *MYC2-MYB* (pLDDT 55.96, FigS2) by docking ATP and acetyl-CoA to predicted structures of *MYC2-COI1* and *MYC2-MYB* complexes respectively.

The best docked pose for ATP on the predicted *MYC2-COI1* complex (Fig 4a, b) showed adenine and tetrahydrofuran groups on ATP were docked inside a *CO11* pocket, with three arginine residues R65, R68, R69 on *COI1* forming hydrogen bonds with four oxygen groups on ATP (indicated by dashed arrows in Fig 4b). R65 is likely an arginine finger as the docked pose of ATP indicates it forms two hydrogen bonds with the γ-phosphate of ATP Fig 4b. An arginine finger is a highly conserved arginine residue found in many enzymes that forms contacts with the γ-phosphate of ATPs/GTPs catalysing the cleavage of the γ-phosphate ^28‒30^. Docking also indicated ATP’s γ-phosphate is solvent accessible and forms van der Waals contacts with threonine (T19), methionine (M16), alanine (A17) and an adjacent hydrophobic glycine (G18) on *MYC2* (Fig 4b). Considering serine, threonine, and tyrosine are the most frequently phosphorylated amino acids in eukaryotes suggests that *COI1* likely phosphorylates *MYC2* via phosphorylation of T19.

**Fig 4:**
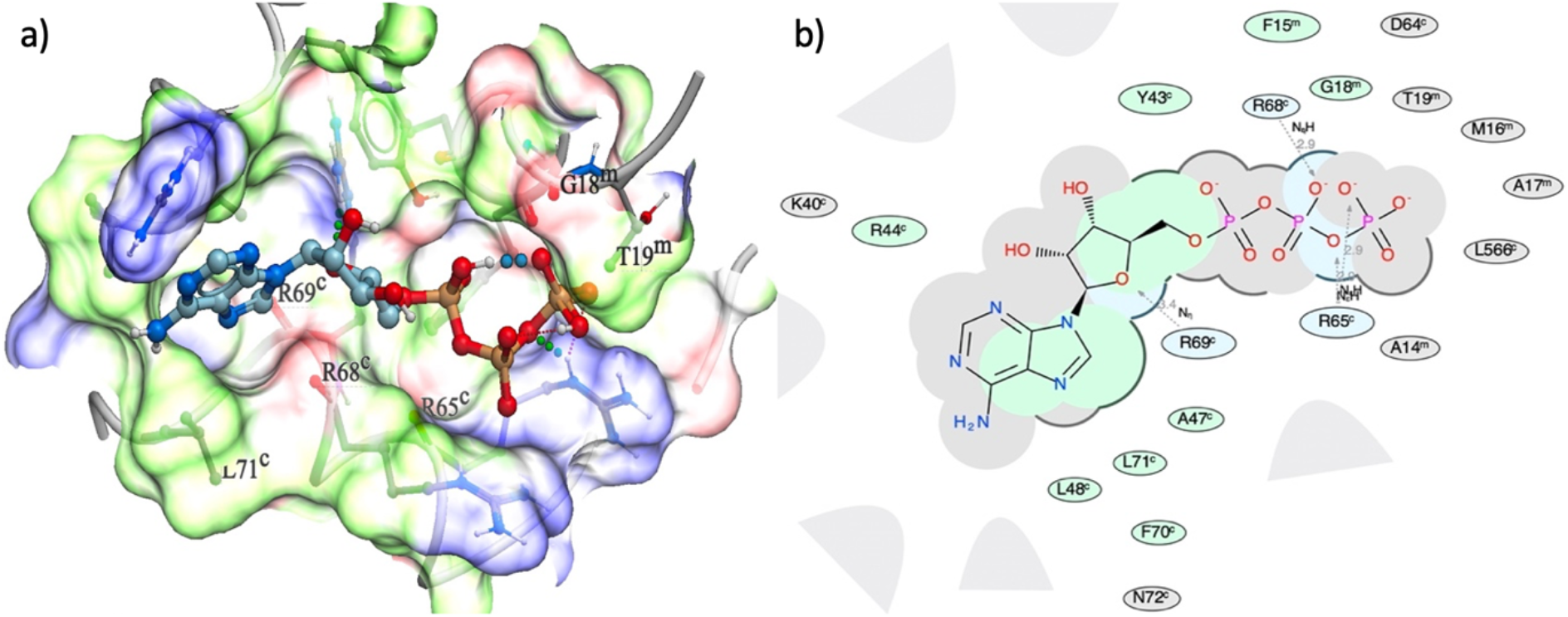
***a)*** ATP docked on a predicted *MYC2-COI1* complex. The binding pocket surfaces are coloured based on the binding properties of the surfaces. White coloured surfaces are potentially aromatic lipophilic, green coloured surfaces are non-aromatic lipophilic, red coloured surfaces are potential hydrogen bond acceptors and blue coloured surfaces potential hydrogen bond donors. Phosphate groups on ATP are coloured brown, nitrogens blue and carbons grey. All phosphates on the docked ATP molecule are exposed towards potential hydrogen bond donors at the opening of the binding pocket (blue coloured surfaces). ***b)*** 2D ligand interaction diagram of ATP interactions with amino acid residues in the binding site. The size of an amino acid label and the distance from ATP represents the interacting area and proximity of the residue to ATP. Grey parabolas and broken thick lines indicate solvent accessible (open) regions, hydrophobic regions are coloured green, blue regions are hydrogen bond accepting and grey regions represent van der Waals contacts. On the predicted *MYC2-COI1* complex, ATP’s adenine and tetrahydrofuran were docked on *COI1*, with the γ - phosphate interacting with residues T19^m^, M16^m^, A17^m^ on MYC2. Residues R65^c^, R68^c^, R69^c^ on *COI1* form hydrogen bonds with four oxygen groups on ATP (dashed arrows).

The best docked pose for acetyl-CoA on the predicted *MYC2-MYB* complex (Fig 5a, b) shows acetyl-CoA binds to *MYC2* likely via hydrogen bonding of acetyl-CoA’s adenine with serine (S150) (Fig 5b). Glutamate (E140 & E155), aspartate (D154) and histidine (H133) were bound to acetyl-CoA’s hydroxyl and phosphate groups (Fig 5b). The predicted binding pose places acetyl-CoA’s acetyl group (^-^COCH3) close to *MYB*’s histidine (K105) (purple residue on Fig 5a, bold green circled residue K105 on Fig 5b) and aspartate (D101) (red residue on Fig 5a, bold green circled residue D101 on Fig 5b). Acetylation of lysine is a well-established regulation process in eukaryotes, known to occur via the transfer of acetyl groups from acetyl-CoA to lysine’s*ε*-amino group ^31^. Inspection of the pocket surface surrounding the acetyl group (Fig 5a) indicates only two residues aspartate and histidine (D101 and K105) can potentially act as a hydrogen bond acceptor (red coloured pocket surface) and hydrogen bond donor (blue coloured pocket surface) respectively with acetyl group. This suggests D101 may act to deprotonate the K105’s*ε*-amine, enabling nucleophilic attack of the acetyl-CoA carbonyl. Similar mechanisms has been proposed previously as likely reaction pathways for acetylation by *histone acetyltransferases* ^32,33^.

**Fig 5:**
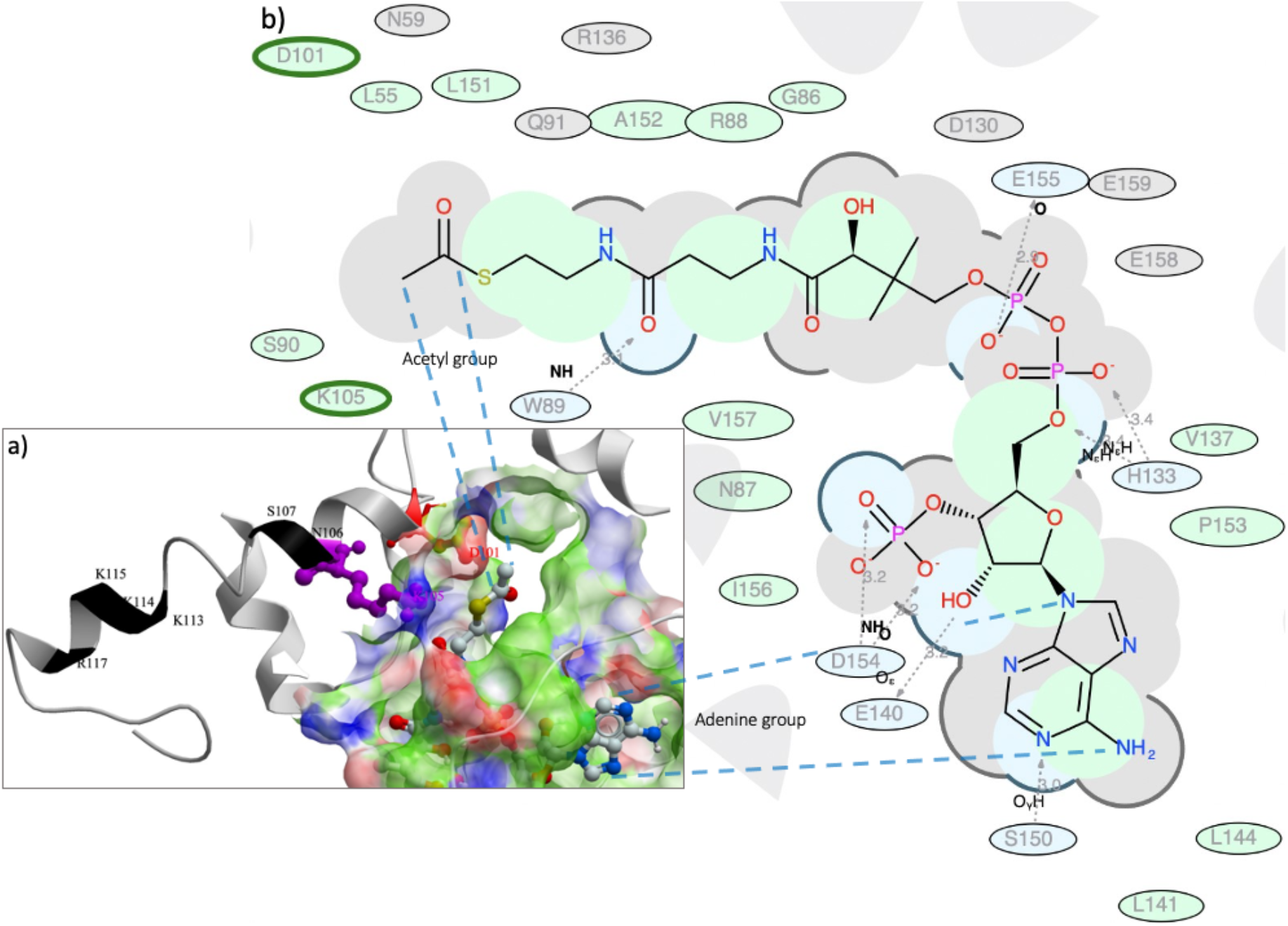
***a)*** Structure of *MYC2-MYB* AlphaFold complex with acetyl-CoA docked. Acetyl-CoA binding pocket surfaces are coloured based on the binding properties of the binding pocket surface. White coloured surfaces are potentially aromatic lipophilic, green surfaces are non-aromatic lipophilic, red surfaces are potential hydrogen bond acceptors and blue surfaces potential hydrogen bond donors. ***b)*** 2D ligand interaction diagram illustrating the interactions of acetyl-CoA with amino acid residues at the binding site. The size of an amino acid label and the distance from acetyl-CoA represents the interacting area and proximity of the amino acid to acetyl-CoA respectively. Grey parabolas and broken thick lines denote solvent accessible (open) regions, hydrophobic regions are coloured green, blue regions are hydrogen bond accepting and grey regions represent van der Waals contacts. On the predicted *MYC2-MYB* complex, acetyl-CoA was docked on *MYC2* with its acetyl group in proximity to *MYB’s* lysine (K105) (highlighted in purple in *a* and a bold green circle in ***b*)**. Residues around K105 (labelled in black in **a**) form part of known conserved acetylation sites. The sequence DNEI**KNS**CI**KKK**LR from residue D101 (highlighted in red in *a* and a bold green circle in *b*), to R117 includes acetylation motifs **KN, K*R, K**^**\***^**K, K*N, K**^**\***^**R** and **K**^**\***^**S**, reported in leaves of paper mulberry, tea, and soybean. The * represents any one residue and the first K in a motif is the acetylated lysine.

Plant proteomic studies have shown that acetylation sites have enriched lysine motifs. These include the motifs below, where an * denotes any one residue. Motifs K*R, K*S, KK, K*K, K****K, and KN, which were previously found to be enriched at acetylation sites in paper mulberry (*Broussonetia papyrifera*) ^34^, are present on *MYB* (black labelled residues on Fig 5a) within 10 residues from K105. Motifs enriched with aspartate were also reported in paper mulberry and *Arabidopsis* ^35^, but these were three and one residue away from the acetylated lysine ^34^. In the docking pose, the likely acetylated lysine (K105) has aspartate (D101, red residue on Fig 5a) four residues away from K105. Proteomics analysis in leaves of tea (*Camellia sinensis*) and soybean (*Glycine max)* also reported similar lysine acetylation site motifs K*R, K*S, KK, K*K, K****K and KN ^36^. These motifs are found on a 14 amino acid stretch (residues 101-117) (DNEIKNSCIKKKLR) on *MYB*. These results suggest *MYC2* likely acetylates *MYB*.

## Discussion

George E.P. Box, the 1950’s British industrial statistician is credited with the quote ‘all models are wrong, some are useful’. Current state of the art deep learning models has significantly improved the usefulness of protein structure and language models in research. The astonishing atomic level accuracy of AlphaFold protein structure predictions, where some predictions are accurate to within 1.4 Å, are revolutionising biological research. However, the predicted structural models are static. Accordingly, structure predictions do not account for the dynamic nature of proteins and the associated modifications that may be necessary for proper function, such as formation of homo or heterocomplexes, presence of ions, water, pH, disordered regions, loops, conformational changes, and post-translational modifications. Despite these limitations, results of predicted proteins and protein-protein complexes combined with functional residue annotation and small molecule docking are encouraging and indicate AlphaFold can be used for coarse grained exploration, with potential to uncover novel insights and mechanisms.

Crystal structures in the PDB typically represent the most dominant conformations of proteins determined experimentally but not *in situ* within cells. Given that AlphaFold and DeepFRI are trained on PDB structures, it is likely their predictions reflect this bias. As a result, protein-protein docking simulations using predicted structures may not be preferable as protein-protein interactions typically involve conformational changes. Rather than docking two proteins individually and then docking small molecules to the resulting protein-protein complex, we explored predicting complexes with AlphaFold and then docking small molecules to the best ranked complex structure. We reasoned that predicted complexes would enable better binding site identification and docking of ATP and acetyl-CoA, allowing us to better infer the likely roles of MYC2, COI1, and MYB based on AlphaFold’s ability to accurately predict protein-protein complexes ^10‒13^. Due to limitations on memory on GPUs, we assumed the complexes would involve single monomers of *MYC2* -*COI1* and *MYC2-MYB*. This reduced the total amino acid length and memory requirements for structure predictions. Docking small molecules to predicted structures of complexes rather than docking two proteins reduced the uncertainty in determining the correct interfaces between two docked proteins.

When used to annotate genes in the context of a differential expression study, the use of AlphaFold and DeepFRI models can result in more precise inference of key pathways and gene functions. Differential expression analysis and Gene Set Enrichment Analysis (GSEA) produce enriched gene lists containing hundreds to thousands of genes. However, it is very difficult to study all genes listed as differentially expressed or enriched and it is often desirable to reduce this list down to a practical number of gene targets. The use of protein structure and language models, as demonstrated, provides an alternative approach that, in addition to reducing gene list numbers, avoids GSEA pitfalls such as database composition bias and artificially inflated GO terms from genes involved in multiple pathways ^37^. Once trained, protein structure and language models can be used independently of the databases used in training. Furthermore, validation methods used during model training reduce the influence of over-represented annotations, reducing database biases caused by the overrepresentation of annotations from duplicate entries from model organisms. In addition, the neural networks that underpin these models can predict novel folds and functions that are not present in the databases used for model training. When applied to a set of genes that share functional annotation terms, these techniques have the potential to be more informative and highly accurate compared to single gene laboratory assays.

We were able to infer from a set of differentially expressed genes that *COI1* likely phosphorylates *MYC2*, and *MYC2* likely acetylates *MYB*, with *MYC2* interacting with multiple genes. Our findings are supported by previous research based on extensive laboratory experiments. Furthermore, the genes we identified as likely interacting (*MYC2, MYB* and *COI1*) have been reported to suppress resistance to several pathogens. Plant mutants with abolished expression of *MYC2* have shown increased resistance to *Fusarium oxysporum, Plectosphaerella cucumerina* and *Botrytis cinerea* ^38‒40^. *COI1* and *MYC2* mutants have also been shown to increased resistance to *Fusarium oxysporum* ^41,42^. Mutation-derived resistance is likely due to MYC2 negatively regulating salicylic acid pathways ^43,44^, inferring the interaction between *MYC2* and *COI1* could be a source for engineered disease resistance in bananas. This suggests that AlphaFold and DeepFRI models combined with small molecule docking can be used to accurately identify genes and residues for engineering novel traits such as disease resistance in addition to functional annotation in non-model organisms.

While the use of deep learning models in plant genomics is still in its infancy, the next decade will see a further leap in model accuracy as better and improved models are applied and experimentally validated. This will allow for more accurate functional annotation in non-model organisms, hypothesis formulation and inform the design of laboratory validation experiments.

## Methods

### Protein structure prediction

Protein sequences of 19 differentially expressed genes in response to *Pseudocercospora fijiensis* in Williams and Calcutta 4 banana varieties identified by ^18^ were retrieved from the Banana Genome Hub database ^45^. The protein sequences were modelled with both AlphaFold ^46^ and the RosettaFold ^7^. Models were retrieved and the corresponding pLDDT scores scaled to between 0 and 1. For each protein, we calculated a TM-score between that protein’s AlphaFold and RosettaFold model. AlphaFold and RosettaFold scaled pLDDT scores and TM-scores were statistically compared using the R MASS package. Due to GPU memory constraints, we did not investigate whether individual genes folded into homo or hetero complexes. We assumed that interacting genes formed hetero complexes, with each protein contributing a monomer. We thus predicted structures of *MYC2-COI1* and *MYC2-MYB* complexes for docking experiments using AlphaFold and Tesla T4 GPUs.

### Functional annotation of residues

We applied a graph convolutional network model DeepFRI ^1^ trained on protein structures and their associated GO terms to predict MF GO terms and EC numbers. Grad-CAM maps for site specific residues associated with a specific GO term were plotted using DeepFRI’s inbuilt plotting module. Grad-CAM values along a protein structure model were visualized on structures using Pymol with residues coloured with a divergent red, white, blue gradient.

### Identification of ATP and acetyl-CoA binding pockets and docking

Potential small molecule binding pockets were identified on structures of *MYC2-COI1* and *MYC2-MYB* complexes using the ICMPocketFinder algorithm ^47,48^. We considered only pockets occurring at the interface of *MYC2* with *COI1* or *MYC2* with *MYB* in the predicted complexes. This eliminated the possibility of allosteric binding pockets for ATP or acetyl-CoA that are unlikely to result in phosphorylation or acetylation confounding our analysis. We identified a total of 28 small molecule binding pockets on the *MYC2-COI1* complex and 26 pockets on the *MYC2-MYB* complex (Table S2). One pocket was at the interface of *MYC2-COI1* while seven pockets were at the interface of the *MYC2-MYB* complex. Structure data files (SDF) for ATP (PID 5953) and acetyl-CoA (PID 6302) were downloaded from the PubChem database ^49^ and docked to all predicted interface pockets. For each pocket, conformational samples with full local minimization based on the biased probability Monte Carlo (BPMC) procedure ^50^ were generated with low energy conformations retained. The bound conformation with the lowest global free energy minimum across all pockets in a gene was taken as the most probable binding site and docking pose of ATP or acetyl-CoA.

## Abbreviations

Cryo-EM: Cryogenic electron microscopy
PDB: Protein Data Base
SLP: Surface layer protein
CASP: Critical Assessment of Structure Prediction
GDT: Global Distance Test
CAFA: Critical assessment of functional annotation
CNN: Convolutional Neural Network
GCN: Graph Convolutional Network
DeepFRI: Deep Functional Residue Identification
GO: Gene Ontology
Pfam: Protein family
Grad-CAM: Gradient-weighted Class Activation Map
pLDDT: Predicted Local Distance Difference Test
MF: Molecular function
EC: Enzyme commission
LOX8: Lipoxygenase 8
TIFY3B: protein TIFY 3B
DREB1D: dehydration-responsive element-binding protein 1D
ERD15: Early Responsive to Dehydration 15 protein
IAA27: Auxin response protein 27
IAA30: Auxin response protein 30
RD22: Dehydration responsive protein RD22
MYC2: Transcription factor MYC2
COI1: Coronatine-insensitive protein 1
EIN3: ET-insensitive 3
MYB: MYB family transcription factor
JA: Jasmonate
SA: Salicylic acid
GA: Gibberellins
ABA: Abscisic acid
IAA: Auxin
SDF: Structural data files
GSEA: Gene Set Enrichment Analysis
BPMC: biased probability Monte Carlo

## Availability of data and materials

Genes analysed in this study were published by Rodriquez *et al*., *2020* and retrieved from Banana Genome Hub database ^45^. Supplementary files FS1, FS2, Table-S1 and Table-S2 are included with this manuscript.

## Acknowledgements

We thank Google Cloud Compute for access to GPU compute GCP19980904, and Queensland University of Technology’s eResearch computing infrastructure for access to HPC facilities.

## Author contributions

G.S. and P.V. carried out the analysis, P.V. conceptualised and supervised the study. B.D., P.D., R.H., and J.D. all contributed to writing the manuscript and discussion. The manuscript was written, reviewed, and approved by all authors.

## Figures

**Fig S1:**
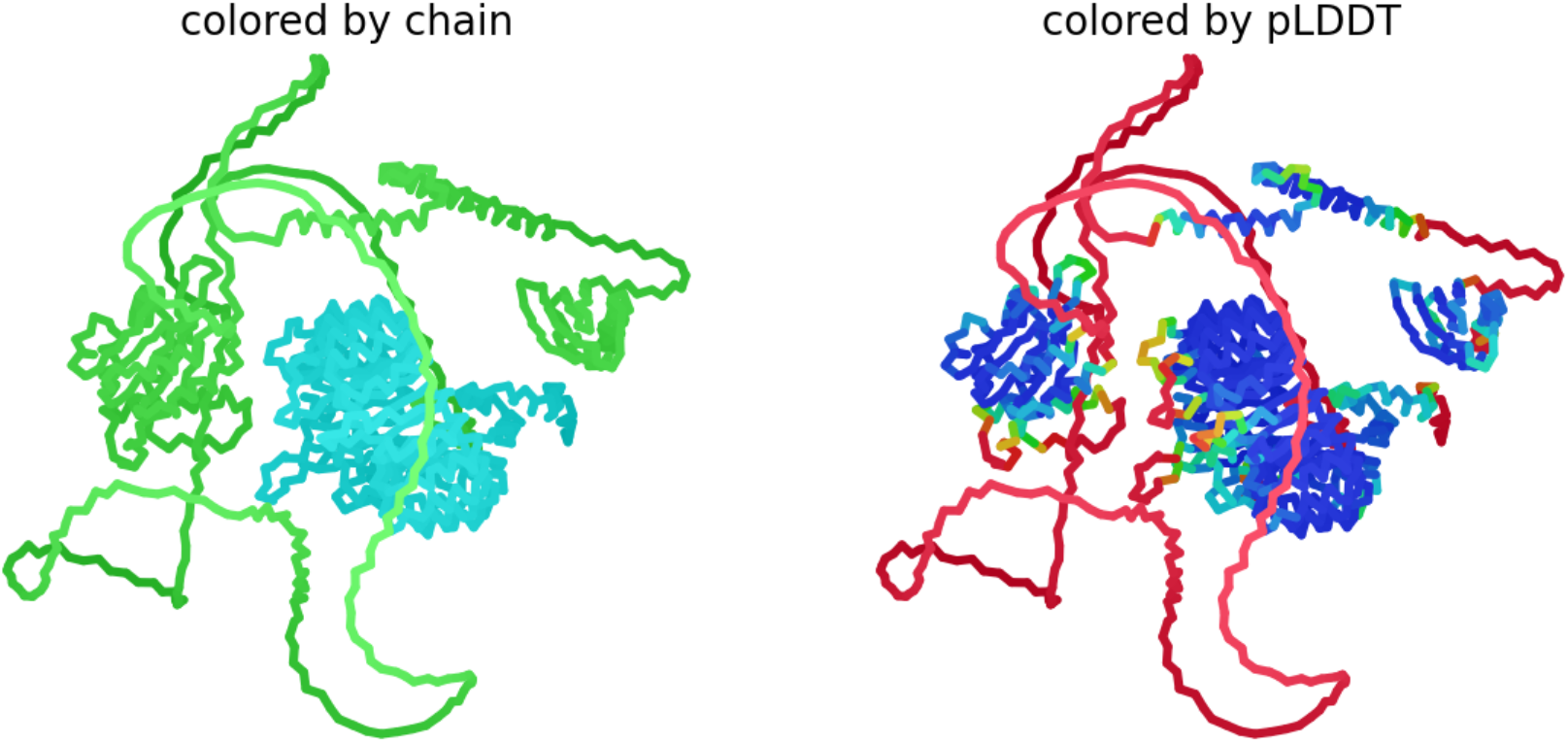
Predicted structure of *MYC2-COI1* complex with a pLDDT of 69.15. Structures are coloured by chain in ***a***, *MYC2* in green, *COI1* in cyan and by pLDDT in ***b***, red being low and blue high confidence regions.

**Fig S2:**
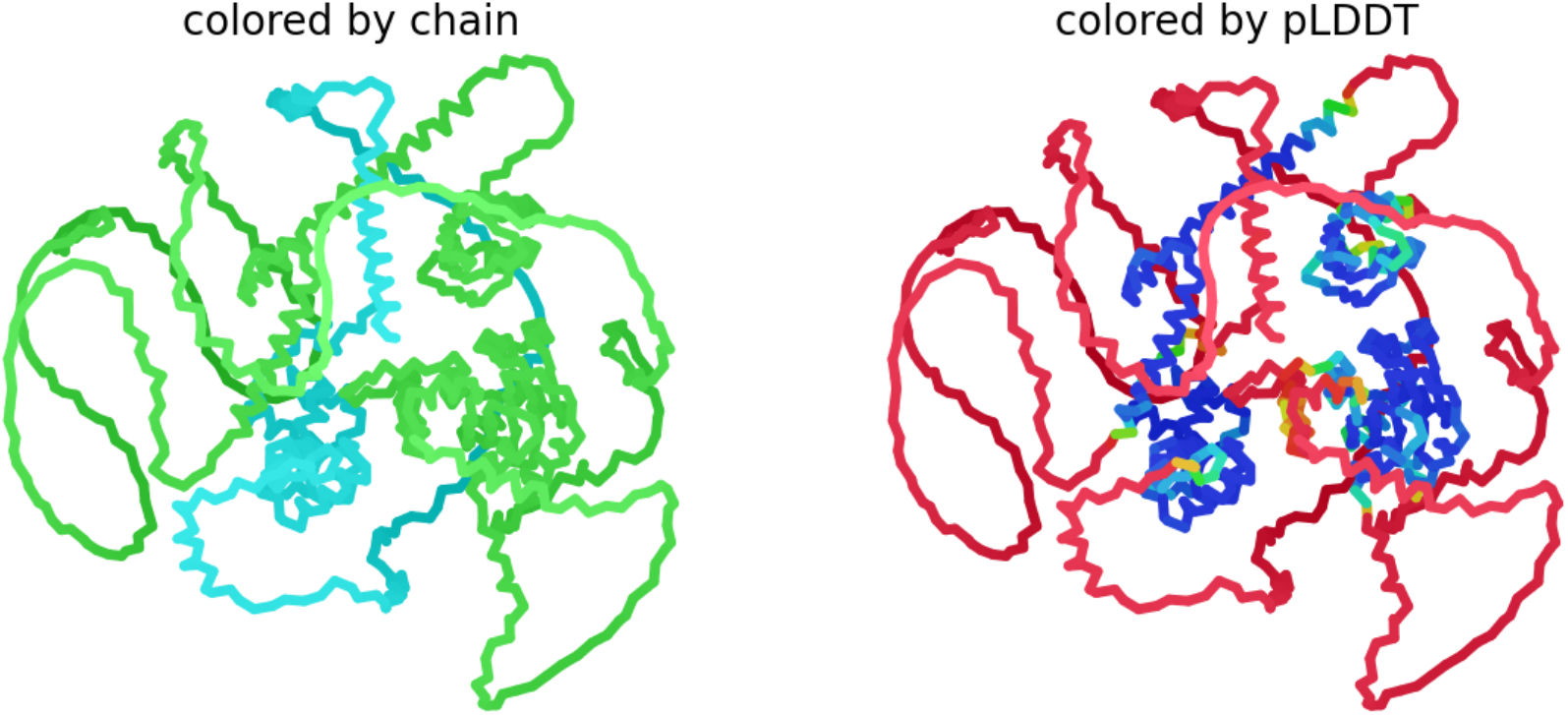
Predicted structure of *MYC2-MYB* complex with a pLDDT of 55.96. Structures are coloured by chain in ***a***, *MYC2* in green, *MYB* in cyan and by pLDDT in ***b***, red being low and blue high confidence regions.

**Table S1:** Deep residue functional annotation with molecular function and EC number assignments for the 19 differentially expressed genes reported by Rodriquez *et al*., *2020*.

**Table S2:** Predicted small molecule binding pockets on ***MYC2-COI1*** and ***MYC2-MYB*** complexes and their pocket properties. Pocket volume (A°), area (A°), aromaticity (fraction of aromatic residue side chains in pocket), radius (A°) and ATP/acetyl-CoA docking scores.

